# Genome-scale rates of evolutionary change in bacteria

**DOI:** 10.1101/069492

**Authors:** Sebastian Duchêne, Kathryn E. Holt, François-Xavier Weill, Simon Le Hello, Jane Hawkey, David J. Edwards, Mathieu Fourment, Edward C. Holmes

## Abstract

Estimating the rates at which bacterial genomes evolve is critical to understanding major evolutionary and ecological processes such as disease emergence, long-term host-pathogen associations, and short-term transmission patterns. The surge in bacterial genomic data sets provides a new opportunity to estimate these rates and reveal the factors that shape bacterial evolutionary dynamics. For many organisms estimates of evolutionary rate display an inverse association with the time-scale over which the data are sampled. However, this relationship remains unexplored in bacteria due to the difficulty in estimating genome-wide evolutionary rates, which are impacted by the extent of temporal structure in the data and the prevalence of recombination. We collected 36 whole genome sequence data sets from 16 species of bacterial pathogens to systematically estimate and compare their evolutionary rates and assess the extent of temporal structure in the absence of recombination. The majority (28/36) of data sets possessed sufficient clock-like structure to robustly estimate evolutionary rates. However, in some species reliable estimates were not possible even with ^“^ancient DNA^”^ data sampled over many centuries, suggesting that they evolve very slowly or that they display extensive rate variation among lineages. The robustly estimated evolutionary rates spanned several orders of magnitude, from 10^−6^ to 10^−8^ nucleotide substitutions site^-1^ year^-1^. This variation was largely attributable to sampling time, which was strongly negatively associated with estimated evolutionary rates, with this relationship best described by an exponential decay curve. To avoid potential estimation biases such time-dependency should be considered when inferring evolutionary time-scales in bacteria.

## INTRODUCTION

Estimating the rate of molecular evolution is critical for understanding a variety of evolutionary and epidemiological processes. Rates of molecular evolution are the product of the number of mutations that arise per replication event, the frequency of replication events per unit time, and the probability of mutational fixation. These rates are determined by a variety of factors including background mutation rate, the direction and strength of natural selection, generation time and population size. Both generation time and population size have been shown to scale negatively with the substitution rate in a range of organisms (Bromham 2009). For example, because of differences in generation time, spore-forming bacteria evolve more slowly over time than those that do not form spores (Weller and Wu 2015). Similarly, lineages undergoing adaptive evolution are expected to accumulate substitutions more rapidly than those subject to purifying selection (Eyre-Walker and Keightley 2007).

Despite the wealth of sequence data, genome-wide rates of evolutionary change in bacteria are often uncertain. At one end of the spectrum, rates as high as ~10^−5^ nucleotide substitutions site^-1^ year^-1^ have been reported for *Neisseria gonorrhoeae* (Pérez-Losada et al. 2007). In contrast, genome-wide rates of only ~10^−9^ substitutions site^-1^ year-1 have been observed in *Mycobacterium tuberculosis* (Comas et al. 2013). Importantly, however, these estimates are not always readily comparable because they use different methods and sources of data. Furthermore, most previous studies have not investigated the degree of temporal structure (i.e. clock-like behaviour) in the data, such that their reliability is uncertain. The increasing availability of genomic data means that it is now possible to estimate substitution rates with sequences collected over a number of years (i.e. tip-date calibration), not just for rapidly evolving pathogens, such as RNA viruses, but also in some DNA viruses and bacteria (Biek et al. 2015). The rates at which bacteria evolve, the strength of clock-like signal in the data, and the determinants of any rate differences observed have not been investigated. However, these are of importance for both the accurate interpretation of outbreak investigations that may depend on the reliable estimation of the time-scale of putatively linked transmission cases, and for revealing the long-term time-scales over which bacteria have been associated with specific hosts.

Importantly, estimates of evolutionary rate have been shown to scale negatively with their time-scale of measurement in several organisms (Ho et al. 2011), a pattern that has been attributed to the gradual purging of deleterious mutations over time (Ho and Larson 2006; Penny 2005). In the context of phylogenetic analyses, the time-scale of measurement corresponds to the age of the calibration or the sampling time-frame (Molak and Ho 2015; Duchêne et al. 2014). In viruses, natural selection, mutational saturation and substitution model inadequacy have also been shown to contribute to this pattern (Duchêne et al. 2014, 2015b; Ho et al. 2015b). However, the phenomenon of time-dependency of rate estimates remains largely unexplored in bacterial genomes, although there is some empirical evidence that it may also hold in these organisms (Comas et al. 2013).

To provide a comprehensive picture of genomic-scale evolutionary rates in bacteria and their temporal dynamics, particularly the extent of time-dependency in the data, we analysed, using a variety of phylogenetic methods, 36 publically available whole genome SNP (single nucleotide polymorphism) data sets from bacterial pathogens associated with human disease sampled over periods extending over 2000 years.

## RESULTS

### Measuring the extent of clock-like structure in bacterial evolution

SNPs can be introduced into bacterial genomes individually via mutations or in clusters by recombination, a process that may bias demographic inferences and rate estimates (Lapierre et al. 2016). Recombination-associated SNPs were therefore removed from each of the 36 bacterial genome alignments using both Gubbins v1.4.2 (Croucher et al. 2014) and RDP4 (Martin et al. 2015), with the latter incorporating several methods to detect recombinant sequences, which were then removed from the alignments. Accordingly, we discarded all putative recombinant sequences from each alignment as well as an entire data set of *Neisseria meningitidis* in which nearly half of the sequences displayed recombination (Supplementary material). Following the removal of recombinant sequences we inferred maximum likelihood phylogenetic trees for each data set under a GTR+Γ substitution model. We then used these trees to perform a regression of root-to-tip genetic distance against year of sampling to assess the degree of temporal structure in each data set. Under clock-like evolution there should be a linear relationship between sampling time (year) and the expected number of nucleotide substitutions along the tree. The slope of the line corresponds to the substitution rate, although this estimate is statistically invalid because data points may not be phylogenetically independent. The extent to which the points deviate from the regression line reflects the amount of among-lineage rate variation, measured using the determination coefficient, R^2^(Rambaut et al. 2016).

This analysis revealed large differences in the degree of clock-like behaviour, with 22 of the 35 data sets having R^2^ values of less than 0.5, suggesting weak clock-like behaviour, and eight with R^2^ values of 0.7 or higher, suggesting stronger clock-like behaviour (Fig. 1). A data set of *Staphylococcus aureus* multi-locus sequence type ST239 had the highest R^2^, at 0.96, with similar values observed in *Vibrio cholerae* (0.92), *Shigella sonnei* (0.89) and *Shigella dysenteriae* type 1 (0.88). The lowest R^2^ was 1.52 × 10^−3^ for *M. tuberculosis* Lineage 2. The regression slope (rate) for *M. tuberculosis* Lineage 2 and *Yersinia pestis* from the second global pandemic included negative values, suggesting that these data sets are have very low rates, or that there is extensive among-lineage rate variation, to allow reliable rate estimation from these data.

**Figure 1.**
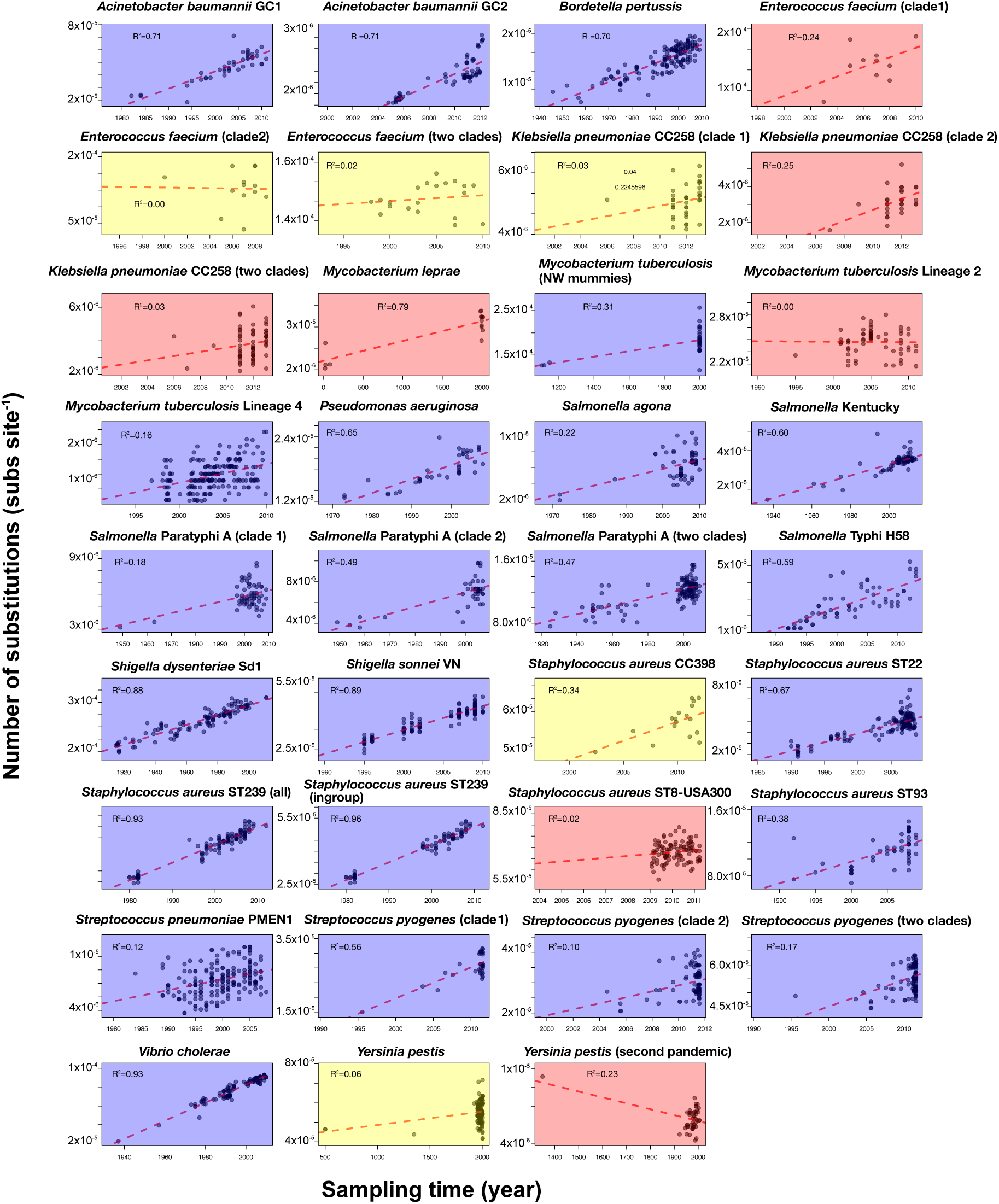
Regressions of the root-to-tip distance (expected nucleotide substitutions site^-1^) as a function of sampling time (year) for 35 bacterial data sets. Each point corresponds to an individual sampled genome (SNP sequence in the alignment), and the red dashed line is the linear regression using least squares; the R^2^ coefficients are also shown. Shading corresponds to degree of temporal structure according to the date-randomisation test in BEAST; blue indicates strong temporal structure, while orange and red indicate moderate and low temporal structure, respective.

We also assessed temporal structure with a Bayesian date-randomisation test (Duchêne et al. 2015a; Ramsden et al. 2009). We used BEAST v1.8 (Drummond et al. 2012) to estimate the rate of evolution according to the optimal molecular clock model selected using the marginal likelihoods. Our comparison of clock models included the strict clock, which assumes that the rate of evolution is constant across lineages, and the uncorrelated lognormal relaxed clock that treats the rate across lineages as a random variable. Most bacterial data sets favoured a relaxed molecular clock, indicating there is evidence for rate variation among lineages (Supplementary material). We then conducted ten date-randomisations replicates per data set, which is sufficient to assess temporal structure in bacterial data (Murray et al. 2015). We classified the degree of temporal structure in each data set according to the number of randomisations whose 95% highest posterior density (HPD) overlapped with that obtained using the correct tip-dates. This analysis suggested that 28 data sets had strong to moderate temporal signal (defined as ≤5 randomisations with 95% highest posterior density (HPD) overlapping with 95% HPDs estimated with the correct tip-dates) (Fig. 2 and Supplementary material). Only one (*M. leprae*) of the 15 bacterial species analysed (excluding *N. meningitidis* which was already discarded because of the large number of recombinant sequences) had no data sets displaying moderate to strong temporal signal. However, five of eight species represented by two or more data sets displayed a mix of both weak signal and strong-moderate temporal signal (Fig. 2), suggesting that a lack of temporal signal may be a property of the individual data sets rather than a true species effect.

**Figure 2.**
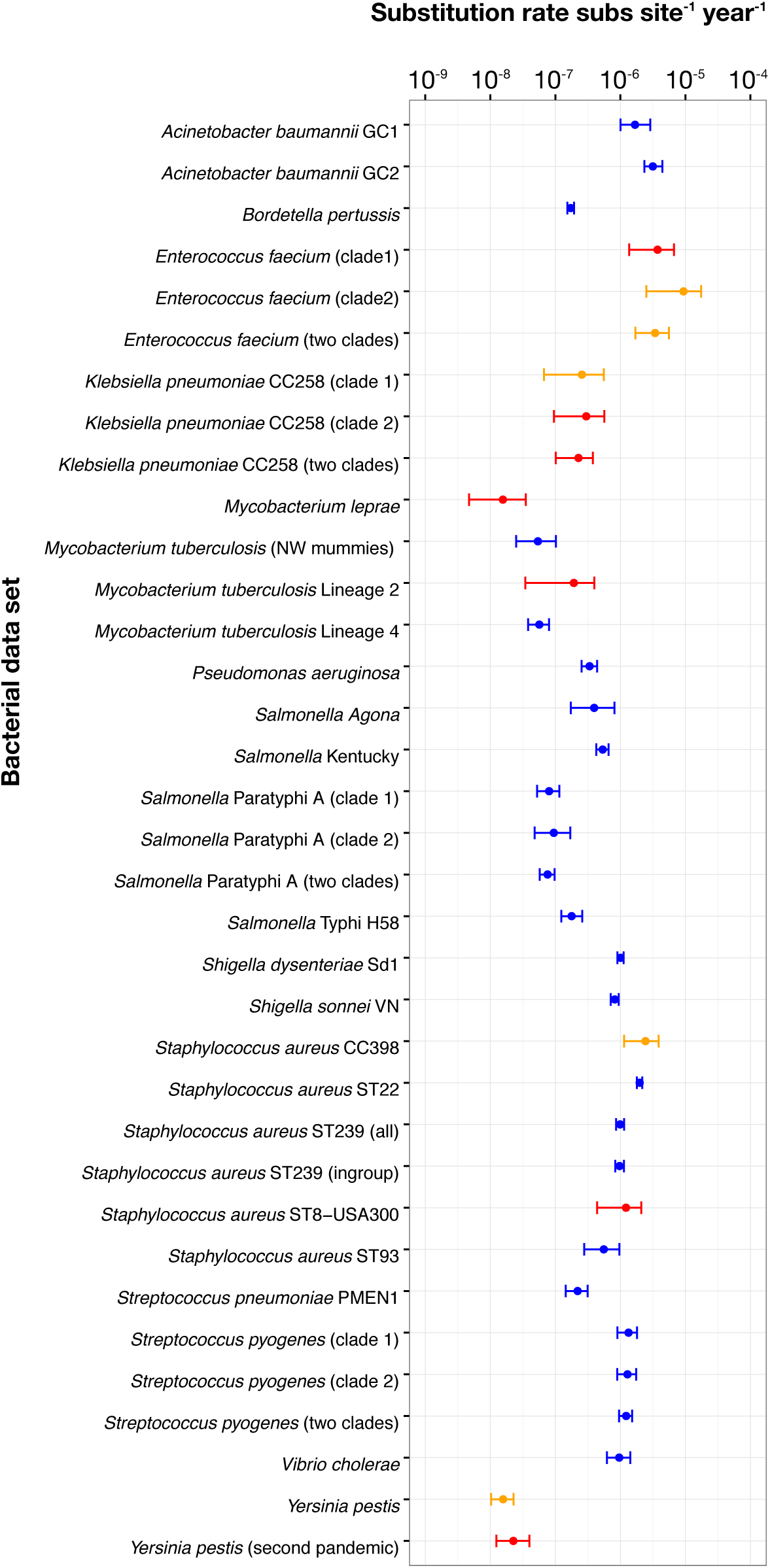
Bayesian estimates of genome-scale nucleotide substitution rates for all bacterial data sets. The axis for the nucleotide substitution rate is shown in log_10_ scale. Points represent the mean rate estimate, and error bars correspond to the 95% HPD values. Colours indicate the degree of temporal structure according to the date-randomisation test as indicated in Fig. 1.

### The range of genome-wide rates of evolutionary change in bacteria

The highest mean genome-wide rate estimate for the data sets with temporal structure was 9.35 × 10^−6^ substitutions site^-1^ year^-1^ (95% HPD: 2.50 × 10^−6^ – 1.74 × 10^−5^) for a vancomycin-resistant *Enterococcus faecium* (VRE) data set sampled over 10 years. Similarly high rates were observed in *Acinetobacter baumannii* Global Clone 2 (GC2) sampled over 7 years and with a mean rate of 3.15 × 10^−6^ substitutions site^-1^ year^-1^ (95% HPD: 2.34 × 10^−6^ – 4.44 × 10^−6^), and *Staphylococcus aureus* clonal complex 398 (CC398) sampled over 9.5 years and with a mean rate of 2.43 × 10^−6^ substitutions site^-1^ year^-1^ (95% HPD: 1.14 × 10^−6^ – 3.98 × 10 ^6^). The lowest estimate was for *Y. pestis* with samples collected over ~1500 years which resulted in a mean rate of 1.57 × 10^−8^ substitutions site^-1^ year^-1^ (95% HPD: 1.03 × 10^−8^ – 2.27 × 10^−8^), although this data set had only moderate temporal structure. Other data sets with low rates and strong temporal structure were: *M. tuberculosis*, with samples collected over ~900 years and a mean rate of 5.39 × 10^−8^ substitutions site^-1^ year^-1^ (95% HPD: 2.49 × 10^−8^ –1.02 × 10^−7^); *M. tuberculosis* Lineage 4 with samples collected over 13 years and a mean rate of 5.67 × 10^−8^ substitutions site^-1^ year^-1^ (95% HPD: 3.80 × 10^−8^ – 8.02 × 10^−8^); and three *Salmonella enterica* serovar Paratyphi A data sets, with sampling times of 58 – 84 years and mean rates ranging from 7.60 × 10^−8^ to 9.47 × 10^−8^ substitutions site^-1^ year^-1^ (Fig. 2 and Supplementary material).

There was a large variation in rate estimates for some closely related bacteria. Our analyses included several data sets of different serovars of *Salmonella enterica*: Kentucky, Agona, Typhi, and Paratyphi A. Interestingly, the rate estimates for the human-restricted serovars Typhi and Paratyphi A (the agents of typhoid fever) (mean rates ranging from 1.78 × 10^−7^ to 8.02 × 10^−8^ substitutions site^-1^ year^-1^) were consistently lower than those estimated for the host-generalist serovars Kentucky and Agona (5.34 – 3.95 × 10^−7^ substitutions site^-1^ year-^1^) over similar sampling times. The rate estimates for *Staphylococcus aureus* were also highly variable between lineages, which included CC398, type USA300 and multi-locus sequence types ST22, ST93, and ST239. The estimated mean rate for the livestock-associated CC398 was 2.43 × 10-^6^ substitutions site-^1^year^-1^ (95% HPD: 1.14 × 10^−6^ – 3.86 × 10^−6^), the highest for this species. In contrast, the estimated mean rate for ST93 was 5.55 × 10^−7^substitutions site^-1^ year^-1^ (95% HPD: 2.77 × 10^−7^ – 9.60 × 10^−7^), nearly five times lower (Fig. 2).

We also investigated whether rate estimates based on root-to-tip regression were consistently biased compared to those obtained using the Bayesian approach in BEAST (Fig. 3). If the estimates from the two methods are the same, then they should fall along the line y=x when plotted against each other. Notably, most points fell above the regression line, indicating that the mean Bayesian estimates (y-axis) were higher than those from the regression (x-axis). A likely cause of this pattern is that deep branches in the phylogenetic trees are over-represented in the regression method. If substitution rates do indeed vary in a time-dependent manner (Duchêne, Ho, et al. 2015; see below), such that higher rates are observed toward the present, then rate estimates obtained using regression would be expected to exhibit a downward bias, as observed in Fig. 3. The exceptions were three *Klebsiella pneumoniae* data sets (CC258), only one of which had sufficient temporal structure for reliable estimation, which yielded a mean rate estimate using BEAST of 2.99 × 10^−7^substitutions site^-1^year^-1^ (95% HPD: 9.51 × 10^−8^– 5.65 × 10^−7^), while that using regression was 9.24 ×10^−6^ substitutions site^-1^ year^-1^ (95% confidence interval, CI: 1.03 × 10^−7^ – 5.07 × 10^−7^); and *Bordetella pertussis*, with a mean rate of 1.74 × 10^−7^ substitutions site^-1^ year^-1^ using BEAST (95% HPD: 1.53 × 10^−7^ – 1.94 × 10^−7^), and of 2.15 × 10^−7^ substitutions site^-1^ year^-1^ (95% CI: 2.24 × 10^−7^ – 2.78 × 10^−7^) using regression.

**Figure 3.**
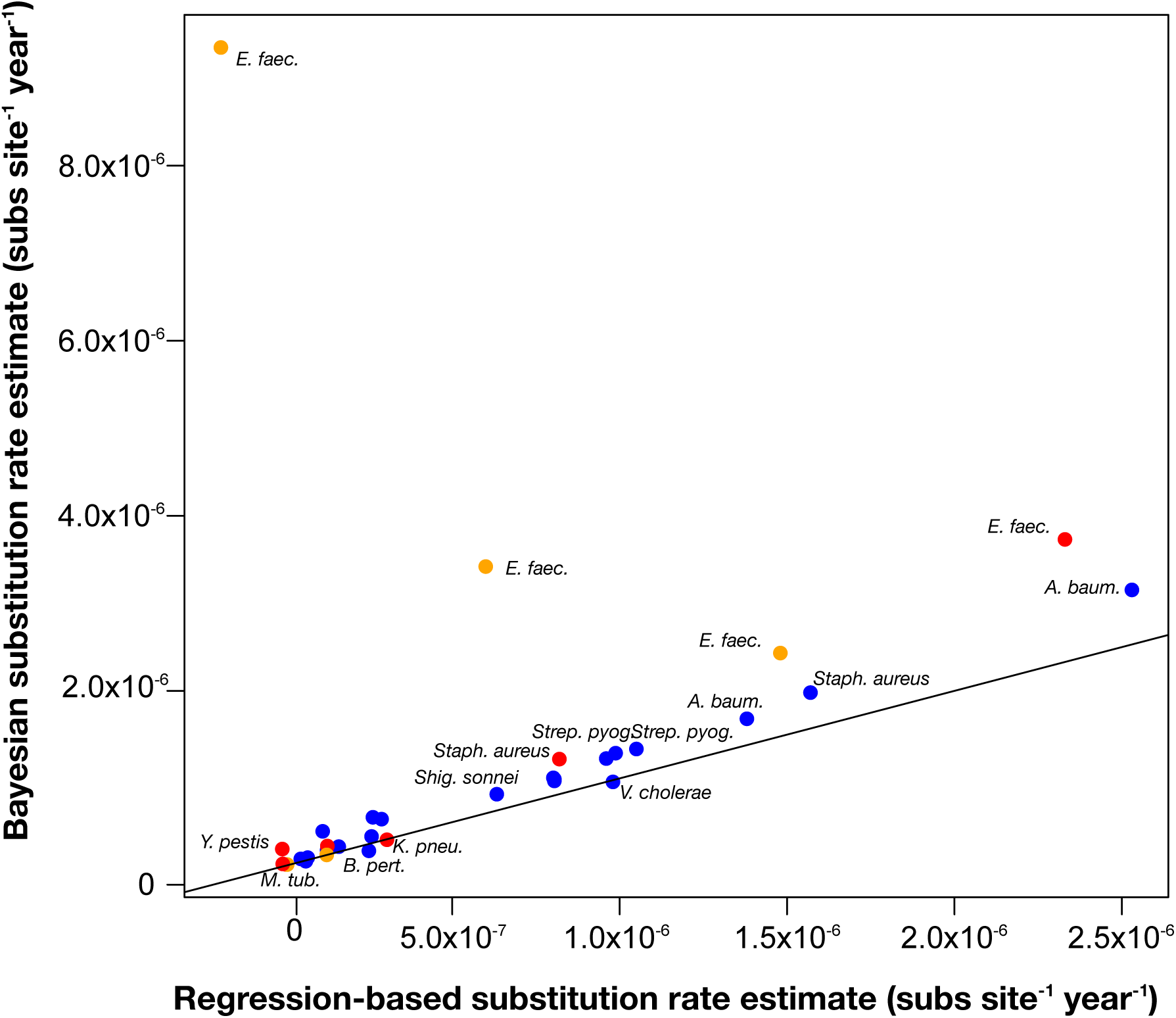
Estimates of genome-scale nucleotide substitution rates using a Bayesian method (BEAST) compared to those estimated via root-to-tip regression. The line represents y=x. Points that fall on the line correspond to data sets for which the mean Bayesian estimates closely match those from the regression. Points above and below the line are data sets for which the Bayesian estimate is higher or lower than that from the regression, respectively. The shading corresponds to the degree of temporal structure according to the date-randomisation test as indicated in Fig. 1.

### Modelling time-dependent rates of evolution

For those data sets with strong to moderate temporal structure, we investigated the relationship between evolutionary rate and sampling time, considered as the time span between the youngest and oldest samples in each data set. To this end we fitted a linear regression for the rate estimates as a function of sampling time on a log_10_ scale, which revealed a significant negative association between rate and time with slope = −0.701 (95% CI: −1.04 to −0.37 and *P* = 0.0003; Fig. 4). However, the linear regression method does not provide a realistic description of such time dependency, which has been suggested to follow an exponential decay curve (Ho et al. 2011; Penny 2005). We therefore modelled the relationship between rate and time using the function *r*=*a*/*T^b^*, in which *r* and *T* correspond to the evolutionary rate and sampling time in log_10_ scale, respectively, and parameters *a* and *b* control the asymptote and rate of decay in the function. This parameterisation is similar to that proposed by O’Fallon (2010). Accordingly, our optimisation of the decay function had the parameterisation *a*=-5.90 (95% CI: −6.39 to −5.77) and *b*=-0.17 (95% CI: −0.26 to −0.13). Importantly, the residual errors were normally distributed around the fitted curve (Supplementary material).

**Figure 4.**
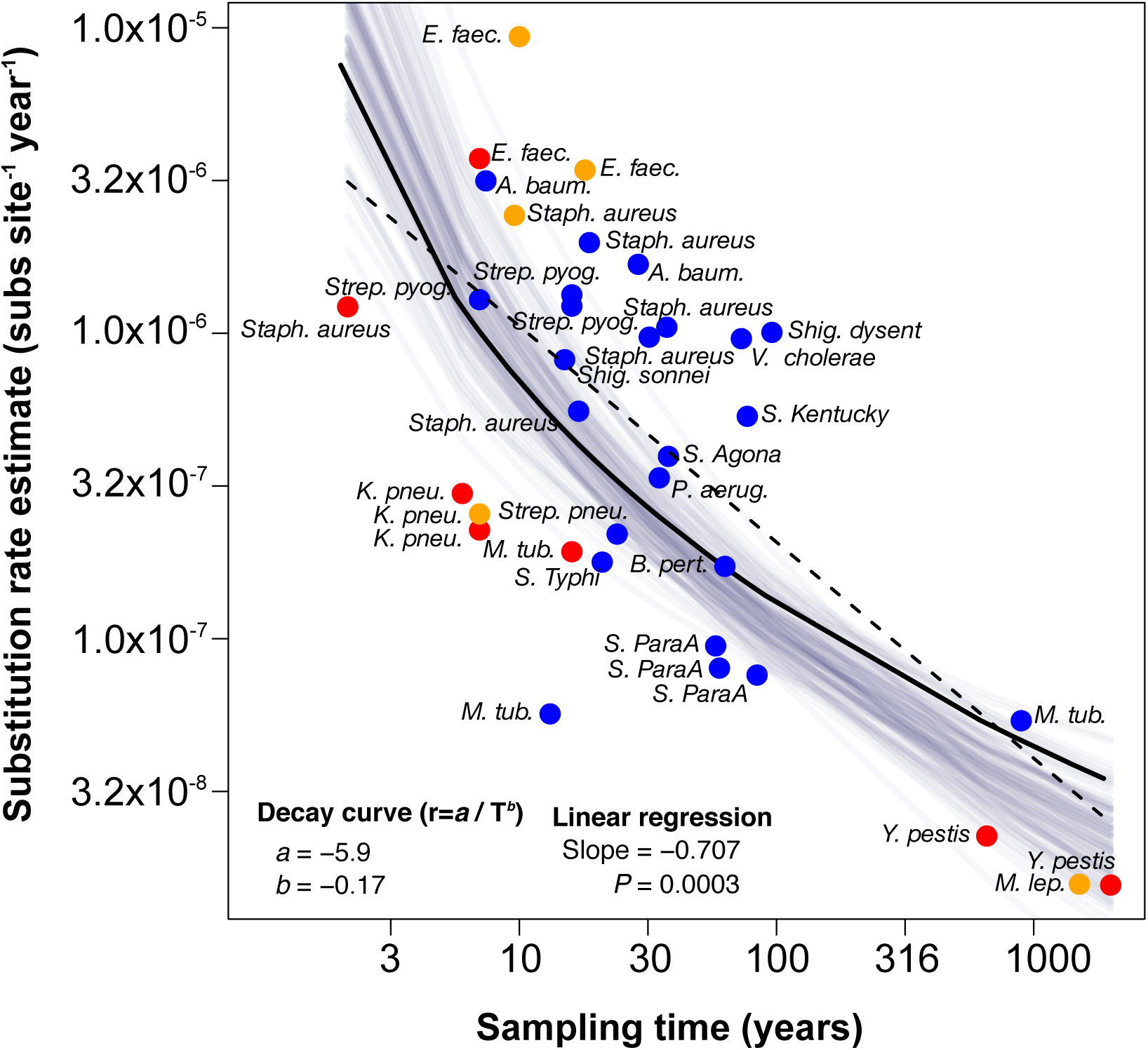
Estimates of genome-wide nucleotide substitution rates in human-associated bacterial pathogens as a function of sampling time in years. The axes are shown on a log_10_ scale. The shading corresponds to the degree of temporal structure according to the date-randomisation test; blue indicates strong temporal structure, while orange and red indicate moderate and low temporal structure, respectively. The dashed line corresponds to the linear regression, while the solid line corresponds to the decay curve (both fitted using only the points with strong and moderate temporal structure). The grey lines represent 100 bootstrap replicates of the decay curve, and thus represent the uncertainty in the decay function.

## DISCUSSION

We have performed the largest comparative and systematic study of evolutionary rates in bacteria to date, providing an important resource for future studies of bacterial evolution. This analysis revealed an approximately two order of magnitude range of evolutionary rates for those bacterial data sets in which there was sufficient temporal structure for rate estimation. For the most rapidly evolving bacteria analysed here, such as *E. faecium*, *S. aureus*, and *A. baumannii*, we estimated genome-wide rates in the order of 10^−6^ substitutions site^-1^ year^-1^. Not only are these rates within the range of those previously estimated for rapidly evolving bacteria (Ward et al. 2014; Holt et al. 2012) but, more strikingly, they are also similar to those of slowly evolving DNA viruses (Duchêne et al. 2014; Firth et al. 2010). However, even our highest rate estimates are lower than some reported previously, such as those for *Helicobacter pylori* (Kennemann et al. 2011) and *N. gonorrhoeae* (Pérez-Losada et al. 2007), at ~10^−5^ and 10^−4^ substitutions site^-1^ year^-1^, respectively. As these species also experience high rates of recombination (Yahara et al. 2016; Maixner et al. 2016), which can disrupt temporal signal, we suggest that these unusually high estimates be treated with caution until they are verified. For example, our *N. meningitidis* data set exhibited extensive recombination, precluding standard phylogenetic analyses.

The lowest rate estimates obtained here, at ~10^−8^ substitutions site^-1^ year-^1^, were also comparable to previous studies of slowly evolving bacteria, notably *M. tuberculosis* (Kay et al. 2015). That even data sets from slowly evolving bacteria such as *M. tuberculosis*, that exhibit substitution rates in the order of 10^−8^ substitutions site^-1^ year^-1^, possessed moderate temporal signal underscores the power of genome-scale data to estimate evolutionary dynamics. Rate estimates lower than about 1.5 × 10^−8^ substitutions site^-1^ year^-1^ could not be validated using our methods and hence may require data sampled over far longer time periods. However, it was also noteworthy that the *M. leprae* and one of the *Y. pestis* data sets did not have sufficient temporal structure for rate estimation despite the availability of ancient DNA. This is consistent with previous studies that have reported no temporal structure in data sets of *Y. pestis* with sampling time ranges of ~290 (Bos et al. 2016) and ~650 years (Wagner et al. 2014), and of *M. leprae* with a sampling range of almost 2000 years (Schuenemann et al. 2013). Interestingly, a recent analysis of Bronze Age strains of *Y. pestis* is consistent with a much higher substitution rate in this bacterium, at ~10^−7^ substitutions site^-1^ year^-1^ (Rasmussen et al. 2015), in contrast to the data presented here and previously (Morelli et al. 2010; Cui et al. 2013). Why 298 these data sets differ so fundamentally in estimated substitution rate is clearly an important area for future study.

Nearly all bacterial species investigated here displayed genome evolution that was measurable over a period of 10-100 years. Data sets with sampling times spanning fewer than 10 years were largely unreliable. Importantly, for several species the strength of temporal signal varied substantially between data sets, and a lack of temporal signal in one data set could not be taken as a general indication of overall lack of signal for the species.

Notably, our analysis reveals that evolutionary rates in bacteria display a strong negative relationship with sampling time, such that their rates can be considered time-dependent (Ho et al. 2011). This observation is of particular importance in the context of the analysis of genome sequences taken from individual disease outbreaks or transmission chains (Didelot et al. 2012), as the evolutionary rates measured over these very short time-scales will likely contain deleterious mutations yet to be purged by purifying selection and substitutional saturation is increasingly apparent over time. As such, these estimates are expected to be much higher than those for samples obtained over longer time-scales. For example, based on our data (Figure 4), we would predict the evolutionary rate estimated for a given bacterial species or clade over a sampling frame of 10 years to be more than an order of magnitude higher than that estimated for the same bacteria sampled over a period of 100 years. Conversely, the longer-term (and lower) evolutionary rates of the type inferred here would tend to over-estimate the time-scale of bacterial transmission when applied to outbreak data. To this end, the relationship between evolutionary rate and sampling time can be used to assess the extent to which extrapolating rates of evolution over different time-scales will lead to a bias in estimates of divergence times.

Strikingly, in some cases the time-dependent pattern appears to hold within closely related bacterial lineages, such as our *A. baumanii* and *S. enterica* Paratyphi A data sets. However, it is notable that both reliable rate estimates for *M. tuberculosis*, estimated over sampling frames of 15 years and 895 years, were nearly identical. *M. tuberculosis* are notoriously slow growing bacteria whose generation time is orders of magnitude slower than the other bacteria analysed here. It is likely that other factors such as genome size and G+C (guanine and cytosine) content also contribute to rate variation between bacteria (Rocha et al. 2006). Given the time dependency of rate estimates we observed, we suggest future studies of bacterial evolutionary dynamics would best addressed by studies that compare multiple independent replicate sample sets for each species, collected over matched time spans. Overall, our analyses show that genome evolution can now be reliably measured in bacteria and establish a genomic framework for understanding long-term evolutionary dynamics in bacteria.

## MATERIALS AND METHODS

### Data collection

We collected 35 whole genome nucleotide SNP alignments from previously published studies (Schuenemann et al. 2013; Wagner et al. 2014; Eldholm et al. 2015; Bratcher et al. 2014; Zhou et al. 2013; Bart et al. 2014; Holt et al. 2008; Merker et al. 2015; Croucher et al. 2011; Marvig et al. 2013; Zhou et al. 2014; Stinear et al. 2014; Gaiarsa et al. 2015; Holt et al. 2013; Devault et al. 2014; Baines et al. 2015; Uhlemann et al. 2014; Davies et al. 2015; Holt et al. 2016; Holden et al. 2013; Howden et al. 2013; Schultz et al. 2016; Ward et al. 2014; Bos et al. 2014; Njamkepo et al. 2016), and one unpublished data set of *Salmonella enterica* serovar Kentucky (Supplementary material). The *S. enterica* serovar Kentucky data set comprised 88 strains isolated between 1937-2012 (Le Hello et al. 2013). Whole genome sequencing was performed using Illumina HiSeq and analysed as described previously (Holt et al. 2013). Although sequence reads for most other data sets are publically available in the NCBI Short Read Archive (SRA), we obtained the original alignments from the authors wherever possible. With this approach we take advantage of domain knowledge of the original studies for choice of reference sequence and identification of repetitive or horizontally transferred sequences, which are important for the accurate generation of SNP alignments. We removed outgroup taxa and samples that were distantly related to the majority of sequences to limit our analyses to the taxonomic group of interest and to minimise the artificial inflation of among-lineage rate variation due to the presence of very long branches in the phylogenetic trees. For the *N. meningitidis* and *M. tuberculosis* Lineages 2 and 4 data sets we obtained sequence reads from SRA and called SNPs using the RedDog pipeline (as described previously (Schultz et al. 2016); available at https://github.com/katholt/RedDog).

Importantly, we removed all genomic regions with evidence of recombination using Gubbins v1.4.2 (Croucher et al. 2014), and verified these results using RDP4 (Martin et al. 2015). For RDP4 we removed sequences with significant evidence of recombination according to six of the methods implemented in the program; Bootscan (Salminen et al. 1995), Geneconv (Padidam et al. 1999), Maxchi (Smith 1992), Siscan (Gibbs et al. 2000),

RDP (Martin and Rybicki 2000), and 3Seq (Boni et al. 2007). We also visually checked that the alignments were free of additional obviously recombining regions. Our final data set comprised 16 bacterial species from 13 genera, with genome sizes ranging from 1.4 (*Streptococcus pyogenes*) to 6.2 (*Pseudomonas aeruginosa*) Mbp. The data set sizes ranged from 189 (*S. pneumoniae* and *M. tuberculosis* Lineage 4) to 15 (*M. leprae* ancient DNA) sequences, and alignment lengths from 15,394 SNPs (*A. baumannii*) to 402 (*M. tuberculosis* Lineage 4) SNPs.

All the data sets included sampling times associated with each sequence in the form of year of isolation. Individual data sets spanned sampling ranges of approximately 2000 years in the case of *M. leprae*, to 2.5 years in *Staphylococcus aureus* ST8:USA300. Four data sets included ancient DNA samples; *M. leprae* (Schuenemann et al. 2013), two *Yersinia pestis* data sets (Wagner et al. 2014), and one *M. tuberculosis* data set comprising strains sampled from New World mummies, animal strains, and human Lineage 6 (Bos et al. 2014). In *M. leprae*, the oldest sample had an estimated age of approximately 2,000 years (Schuenemann et al. 2013), while in the *Y. pestis* data sets the oldest samples had ages of approximately 600 and 1,500 years (Wagner et al. 2014). The New World mummy *M. tuberculosis* samples had ages of approximately 900 years (Bos et al. 2014). Notably, one of the *Y. pestis* data sets included samples from both contemporary strains and those from the first, second, and third plague pandemics, while the other is a subset that only included sequences from the second pandemic that began with the Black Death and contemporary strains. The remaining data sets had sampling times from a few years to several decades (Supplementary material). All the data sets used here are available online(http://zenodo.org/record/45951#.Vr1M-JN95E4).

### ML analysis and root-to-tip regression

We estimated ML trees using PhyML v3.1 (Guindon et al. 2010) employing the GTR+Γ substitution model with four categories for the Γ distribution of among-site rate heterogeneity and SPR branch-swapping. We did not consider the proportion of invariable sites in the substitution model because the data sets comprised SNPs only, such that all sites are variable. The branch lengths in the ML trees are the expected number of nucleotide substitutions per site; as we are working with SNP alignments, this corresponds to the expected number of substitutions per variable site. To convert the branch lengths into genome-wide distances we multiplied them by the length of the SNP alignment, and divided them by the core genome length that reflects the size of the genome in which SNPs were called (this information was extracted from the original publications). We fitted regression for the root-to-tip distance as a function of sampling time using NELSI v0.21 (Ho et al. 2015a).

### Bayesian analysis and date-randomisations

We estimated rates of evolutionary change using a Bayesian Markov chain Monte Carlo method implemented in BEAST, with a chain length of 10^8^ steps, sampling every 5,000 steps. If the effective sample size of any of the parameters was less than 200, we increased the chain length by 50% and reduced the sampling frequency accordingly. For all data sets we used the GTR+Γ substitution model and the sampling times (tip dates) for calibration. For this analysis we utilized both strict and uncorrelated lognormal molecular clock models and constant-size coalescent and Bayesian Skyline demographic models, resulting in four possible model combinations. We compared the statistical fit of these models by estimating marginal likelihoods via path-sampling (Xie et al. 2011). We report the rate estimates from the model with the highest marginal likelihood.

To validate our Bayesian rate estimates we conducted a date-randomisation test, which consists of repeating the analysis while assigning the sampling times randomly to the sequences (Firth et al. 2010). In all cases we used the best-fit combination of demographic and clock model, described above. We conducted ten randomisation replicates per data set to generate an expectation of the rate estimates in the absence of temporal structure. Two criteria have been proposed to assess temporal structure. One method, known as CR1 (Duchêne et al. 2015a), considers that data do not have temporal structure if the mean rate with the correct sampling times is contained within the credible interval of that from any of the randomisations. CR2 is more conservative; data do not have temporal structure if the credible interval of the estimate with the correct sampling times overlaps with that from any of the randomisations. These criteria have different levels of type I and type II errors (Duchêne et al. 2015a). Here, we considered the proportion of randomised replicates with intervals that overlapped with those obtained using the correct sampling times, following CR2. We arbitrarily determined that data had ‘strong’ temporal structure if the proportion was 0, ‘moderate’ if it was between 0 and 0.5, and ‘low’ if it was less than 0.5. We verified that samples with the same sampling time did not form monophyletic groups in our ML trees. Such pattern can produce false positives in the date-randomisation test (i.e. incorrectly suggesting that a data set has temporal structure) (Duchêne et al. 2015a; Murray et al. 2015).

### Regression analyses of the time dependency of substitution rates

A key element of our study was to determine whether the time-span of sampling was associated with the evolutionary rate estimate. To test for this pattern we conducted a least squares linear regression for the genome-wide rate as a function of sampling time. Importantly, only rate estimates with strong and moderate temporal structure were included in this analysis. We used a log_10_ transformation because our rate estimates span several orders of magnitude. A potential shortcoming of this analysis is that the data do not necessarily represent independent samples because some correspond to closely related lineages. This can be addressed by using phylogenetic independent contrasts, or phylogenetic generalised least squares. However, these methods require a phylogenetic tree of the evolutionary relationships of all data points that cannot be estimated for our data because the sites in different SNP alignments are not necessarily homologous. Instead, we used an approach in which the regression is conducted using a single randomly chosen data point from each species. We repeated this procedure 1,000 times and verified that the range of slope estimates with random subsamples did not include zero, and we report the mean value and the corresponding confidence interval. For our regression of the rate as a function of sampling time we used a test described previously (Duchêne et al. 2014) to verify that the slope estimate was not a statistical artefact that sometimes occurs when fitting a regression for a ratio as a function of its denominator. To model the pattern of a decreasing rate over time more realistically, we fit a decay curve of the form *r*=*a/T^b^* where *r* and *T* correspond to the rate estimate and sampling time in log^10^ scale, respectively. Parameters *a* and *b* were optimised using the Nedler and Mead algorithm (1965). To obtain a confidence interval around the parameter estimates, we conducted 100 bootstrap replicates of the rate estimates and sampling times.

## DATA ACCESS

All data sets used in this study are available online (http://zenodo.org/record/45951#.Vr1M-JN95E4JN95E4).

## ACKNOWLEDGEMENTS

This research was funded by the NHMRC (Australia Fellowship #AF30 to E.C.H., Career Development Fellowship #1061409 to K.E.H.). S.D. was supported by a McKenzie fellowship from the University of Melbourne. The following researchers kindly provided us with sequence alignments: Johannes Krause (*M. leprae*), Alexander Herbig (*M. leprae*), Zhemin Zhou (*Salmonella* Paratyphi A), Simon Harris (*B. pertussis*), Julian Parkhill (*B. pertussis*), Paul McAdam (*M. tuberculosis* Lineage 2, and *S. aureus* CC398), Stephano Garasia (*Klebsiella pneumoniae*), Sarah Baines (*S. aureus* ST239), Anne-Catrin Uhlemann (*S. aureus* ST-USA300), Mark Davies (*S. pyogenes*), Mark Schultz (*A. baumannii*), Melissa Ward (*S. aureus* CC398) and Iñaki Comas (*M. tuberculosis* New World mummies).

## Supplementary material

**Figure S1.** Estimated genomic substitution rates in human-associated bacterial pathogens and tip-date randomisations. Points correspond to the mean estimate and the error bars are the 95% HPD. Data points in black are for the estimates obtained with the correct sampling times, while those in orange were obtained using the tip-date randomisations.

**Figure S2.** Heatmap of statistical support for different combinations of molecular clock and demographic models, estimated using marginal likelihood. Rows correspond to bacterial data sets and are sorted in descending order according to their Bayesian mean genome-wide rate estimate, as shown. The columns represent the different combinations of molecular clock and demographic models; cell colours indicate the ranking of statistical support for each data set; from red (highest marginal likelihood) to light yellow (lowest marginal likelihood).

**Figure S3.** Residual plots for rate as a function of sampling time according to the decay function y=-5.93/x^-0.17^, where y and x correspond to the substitution rate and sampling time in log_10_ scale. (A) shows the residuals as a function of the fitted substitution rate, where each point corresponds to a bacterial data set. (B) is the probability density of the residuals.

**Table S1.** Information on the bacterial data sets used in this study, including parameter estimates and the corresponding references.

